# Environmental influences on the mating system of the common morning glory

**DOI:** 10.1101/138982

**Authors:** D. F. Alvarado-Serrano, S-M. Chang, R. S Baucom

**Affiliations:** Ecology and Evolutionary Biology Department, Univers ity of Michigan; Department of Plant Biology, Univers ity of Georgia

**Keywords:** inbreeding, *Ipomoea purpurea*, mating system, morning glory, selfing

## Abstract

The balance between selfing and outcrossing is a life history trait of major concern with deep evolutionary consequences in mixed mating species. Yet, our current understanding of the proximate and ultimate determinants of species’ mating system is still unsatisfactory and largely theoretical. Indeed, evolutionary biologists are still puzzled by the often dramatic variation of mating strategies within single species. Of particular concern is the extent to which environmental conditions shape patterns of variation and covariation of mating system components within species. Here, we address this concern in the common morning glory (*Ipomoea purpurea*) by taking advantage of an extensive dataset of floral traits, genetic estimates of selfing and inbreeding, and relevant environmental factors compiled for 22 populations of this species distributed along a disparate set of environments along Southeast and Midwest USA. Combining a powerful array of parametric and model-free statistical approaches, we robustly identify a set of natural and anthropogenic environmental factors underlying population-level variation in selfing, inbreeding, and flower morphology. Remarkably, individual mating system components are found to be associated with different environmental factors and only loosely associated with each other, and thus potentially under multiple different selective pressures. These results not only corroborate theoretical expectations of the significant role the environment plays in the local determination of mating systems, but also provide compelling evidence of complex underlying interactions between multiple evolutionary processes.

## INTRODUCTION

Mating systems influence the genetic structure and diversity of populations and thus are a key component of species’ evolutionary dynamics (Darwin, 1876; Charlesworth, 2006). Mating systems directly impact fitness (Agren and Schemske, 1993) and hence the maintenance of populations (Pujol *et al.*, 2009), as well as the ability of populations to respond to selection (Noël *et al.*, 2017). Thus, not surprisingly the diverse number of mating system types among flowering plants have both interested and puzzled evolutionary biologists for centuries. In particular, the ability of many species to produce progeny through both selfing and outcrossing (i.e., mixed mating systems) and the often dramatic variation of selfing and outcrossing rates within single species have challenged simple evolutionary explanations (Goodwillie *et al.*, 2005; Karron *et al.*, 2012). Several theoretical explanations have been put forward to reconcile these patterns with the strong fitness consequences that alternative reproductive modes carry for their bearers (Kalisz, 1989; Goodwillie *et al.*, 2005; Glémin *et al.*, 2006). Yet, the question of how mixed mating systems are maintained and what explains its variability across populations remains open. What is clear, however, is that underlying these patterns are environmentally driven processes.

Multiple environmentally driven processes are undoubtedly acting simultaneously. Their combined effect on the evolution and maintenance of mixed mating systems depends on their influence on the trade-off between reproductive assurance and inbreeding depression avoidance (Lande and Schemske, 1985; Goodwillie *et al.*, 2005; Barrett, 2014). On one hand, local environmental conditions may limit opportunities for outcrossing (e.g., by reducing the abundance of pollinators; Scheper *et al.*, 2013; Cusser *et al.*, 2016, or conspecifics; Sagarin *et al.*, 2006). Under these circumstances plants should benefit from being able to produce offspring through selfing (Goodwillie *et al.*, 2005)—a capability that, if not counterbalanced by other evolutionary forces (Fisher, 1941; Stone *et al.*, 2014), should be favored given the reproductive assurance it confers. Under these conditions alleles that allow selfing should rapidly increase in frequency given the automatic transmission advantage of self-fertilization (i.e., the proportionately higher representation of selfed genes among offspring). On the other hand, if local environmental conditions do not limit outcrossing opportunities, selfing could be detrimental as it increases the chances of inbreeding depression and pollen discounting (i.e., reduction in the opportunities for pollen to contribute to the outcrossing pollen pool; Chang and Rausher, 1998; Harder and Wilson, 1998; Fishman, 2000). Together these environmentally dependent interactions should ultimately determine the specifics of populations’ mating system (Barret and Eckert, 1990).

Disentangling environmental drivers of mating systems is, however, remarkably complicated because environmental factors influence multiple processes. First, local environmental variation conditions the relative fitness consequences of inbreeding and hence, the relative cost of selfing (Armbruster and Reed, 2005; Cheptou and Donohue, 2011). Second, environmental variation determines the opportunities for outcrossing via its influence on both the extent and effectiveness of pollen movement between and within populations (Friedman and Barrett, 2009; Cranmer *et al.*, 2012) and by shaping traits that are involved in pollinator attraction (Totland, 2001; Nicolson and Nepi, 2005). Furthermore, local population sizes and extirpation probability, and hence the effectiveness of selection, are also dependent on local environmental variation (Byers and Waller, 1999). In turn, these environmentally driven interactions are expected to affect a set of diverse evolutionary processes, including i) the sorting of genetic diversity (Glémin *et al.*, 2006), ii) the interaction between multiple disparate selective pressures (Barrett, 2003; Sargent *et al.*, 2006), iii) the relative risk of maladaptive gene flow (Peterson and Kay, 2015), and iv) the relative complexity of underlying genetic architecture (Holtsford and Ellstrand, 1992; Fishman and Stratton, 2004). Further complicating matters, all these multiple interactions should result in conflicting evolutionary pressures acting on different mating components (Ivey and Carr, 2005; Dudley *et al.*, 2007). Thus, under potentially weak underlying genetic correlations between traits (Lankinen *et al.*, 2007; Dudley *et al.*, 2007), congruent responses among mating system components are expected only under specific evolutionary scenarios—such as strong correlated selection (Endler, 1995; Armbruster and Schwaegerle, 1996). Yet, there is currently limited empirical evidence that investigates this expectation or examines the mechanisms that underlie mating system variation in natural populations (Lankinen *et al.*, 2016).

To improve our understanding of mixed mating systems it is vital to first investigate how selfing and inbreeding are impacted by environmental factors across geography. Also needed are studies that investigate how correlations between mating system components may be affected by environmental variation. Investigating these patterns would lead to a better understanding of the mechanistic processes involved in the evolution of mating systems and ultimately answer the question of how similarly do mating system components—such as the selfing rate and inbreeding coefficient—respond to environmental variation. Here we address this research gap in *Ipomoea purpurea*, a major agricultural weed, using a comprehensive sample of populations distributed across a significant portion of the species range. We explore spatial patterns of mating system variation and uncover the environmental factors that may influence mating in this species. Specifically, we investigate i) whether floral traits are strongly correlated with mating system parameters, ii) whether population variation of mating system parameters and their correlation with floral traits is geographically structured, iii) which environmental factors best predict individual variation of mating system parameters, and iv) whether strength of the floral-mating system correlation is influenced by the environment. By addressing these questions, we offer valuable insights into the proximate determinants of mating system variation and their complex interaction, and provide compelling evidence of the complex nature of the selective pressures acting on mating systems.

## METHODS

### Study system

Our study focuses on *I. purpurea*, a climbing annual vine with a wide distribution across both Central and North America (Ennos, 1981; Defelice, 2001). Appreciated in horticulture for its colorful flowers, this species has become a weed of major agricultural concern worldwide (Baucom and Mauricio, 2004; Fang *et al.*, 2013). *I. purpurea* shows geographic variability in both selfing rates (Kuester *et al.*, 2017) and resistance levels to glyphosate—the most commonly used herbicide in the US (Benbrook, 2016)—with some populations exhibiting 100% survival after application of standard recommended doses of glyphosate and other populations exhibiting high susceptibility (Kuester *et al.*, 2015). Previous research on this hermaphroditic species has identified heritable variation in anther-stigma distance (Chang and Rausher, 1998), a trait that has consistently been found to be linked to selfing rate (Holtsford and Ellstrand, 1992; Duncan and Rausher, 2013), as well as a significant association between resistance to glyphosate and outcrossing rates (Kuester *et al.*, 2017). This species is thus particularly suitable for investigating the role of both natural and anthropogenic environmental variation in the maintenance of mixed mating systems. Here, we focused on 22 populations of *I. purpurea* sampled along a disparate set of environments along Southeast and Midwest USA in 2012 (Fig. 1), and addressed the extent to which mating system trait variation and covariation are impacted by natural and anthropogenic environmental factors.

**Figure 1.**
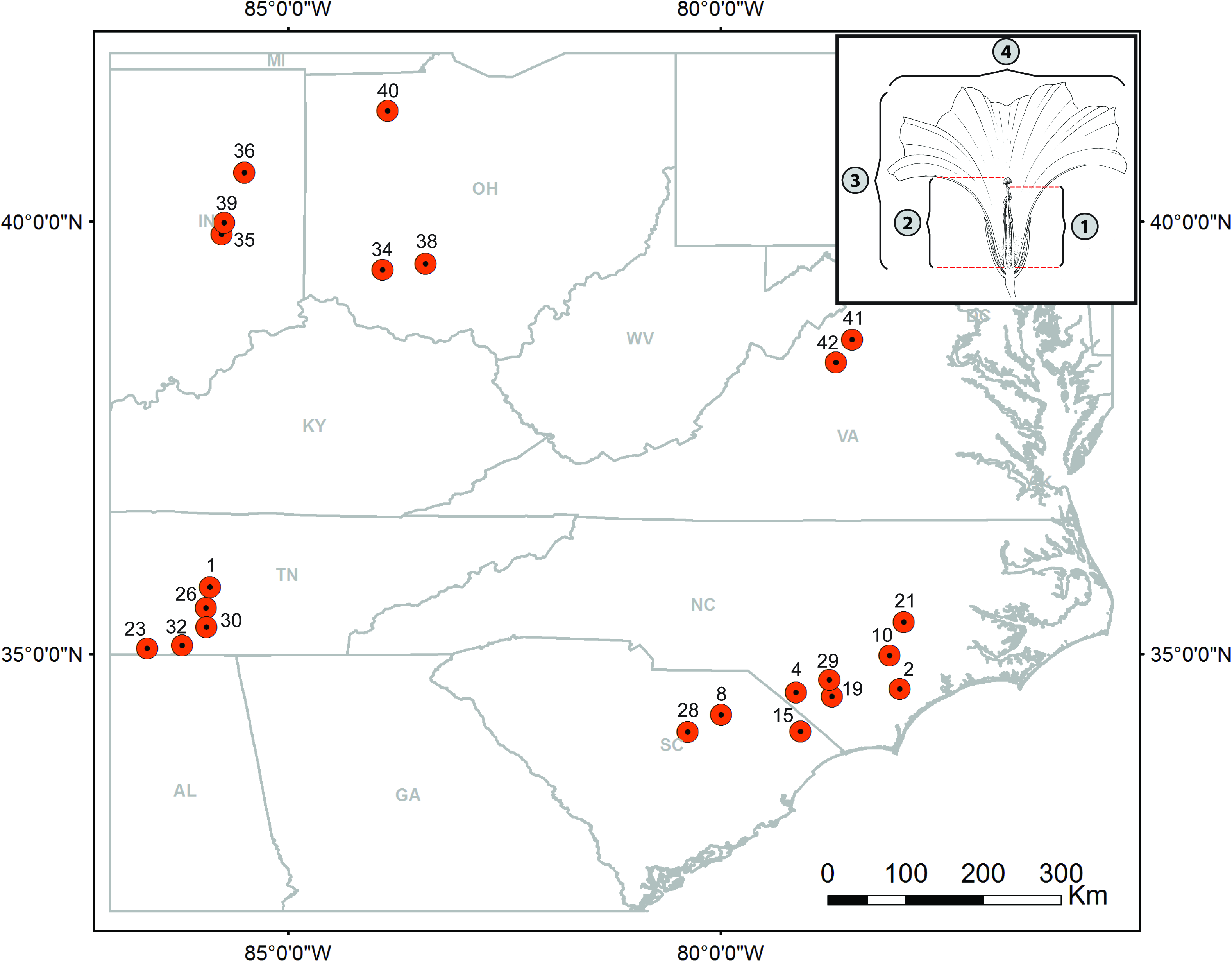
Samples distribution of *I. purpurea* populations with floral measurements and mating system estimates. The four floral measurements taken are shown in upper-right inset: 1) length of the tallest stamen to the top of the anther, 2) height of the pistil to the top of the stigma, 3) corolla length, 4) corolla width.

### Data compilation

To characterize the environments of the 22 sampled populations of *I. purpurea*, we compiled data on a wide range of abiotic factors that could (directly or indirectly) influence mating system variation in this species (Table S1). Given the difficulty of measuring all these environmental factors *in situ*, we chose to use remote sensing and census data. While this decision carries an intrinsic spatial resolution limitation (Holtsford and Ellstrand, 1992), GIS data at moderate to coarse resolutions have been shown to reasonably capture biologically relevant population-level processes (Kerr and Ostrovsky, 2003; Kozak *et al.*, 2008). In addition, because of the previously identified association between resistance to glyphosate and outcrossing rates (Kuester *et al.*, 2017), we included population-level glyphosate resistance estimates— measured as the proportion of surviving individuals after the application of manufacturer’s recommended doses of glyphosate (Kuester *et al.*, 2015), along with county-level estimates of the cumulative amount of glyphosate applied to these populations over the last two decades (years 1992–2012). Unfortunately, no reliable data on bumblebees’ abundance—*I. purpurea*’s primary pollinators (Defelice, 2001)—could be found to be included in our analyses. Our complete environmental dataset included a total of 31 predictor variables (Table S1) with several that were highly correlated with each other. Therefore, we performed a hierarchical agglomerative clustering in R3.3.3 (R Core Team, 2017), using package ClustOfVar (Chavent *et al.*, 2013) to select a non-redundant set of environmental predictors. This analysis clusters variables into statistically homogeneous sets and hence identifies groups of variables that basically bring the same information (Chavent *et al.*, 2012). We chose this analysis because it has the advantage of interpretability over alternative approaches such as principal component analysis (Dormann *et al.*, 2013). By selecting from each resulting cluster the variable less correlated with the other clusters, we retained a set of 8 non-highly-correlated environmental variables (average absolute Pearson’s coefficient = 0.36 [0.01–0.74]).

In addition to the environmental data, a set of four floral measurements were taken over multiple dates in the fall of 2014 from a total of 445 individuals from all 22 populations grown at the Matthaei Botanical Gardens at the University of Michigan (Ann Arbor, MI, USA) (Kuester *et al.*, 2017). Specifically we measured the length of the tallest stamen to the top of the anther (TAL), height of the pistil to the top of the stigma (SL) and the length (CL) as well as width (CW) of the corolla (Fig. 1) on multiple flowers per individual (median number of flowers per individual: 4 [1–25]; median number of individuals per population: 15 [3–23]). In addition, we calculated the difference between the length of the tallest stamen and the height of the pistil, or the anther-stigma distance (ASD). All five floral traits were averaged for each population across flowers, dates, and individuals. Finally, because all four averaged floral measurements (i.e., TAL, SL, CL, CW) were highly correlated with each other (Fig. S1a), we condensed them into a single variable by running a principal component analyses on their covariance matrix after scaling all of them. The retained first principal component, which accounted for 73.77% of the total variance was equitably negatively associated with all four floral measurements included (Fig. S1b), and primarily summarized overall flower size (with lower scores corresponding to bigger flowers). For ease of interpretation, however, populations’ scores on this axis were multiplied by −1 so that flower size increased as PC scores increased. The resulting inverted axis was used as an additional covariate of mating system traits in subsequent analyses.

We quantified mating system estimates for the 22 populations with floral data using individuals genotyped at 15 microsatellite loci previously developed for *I. purpurea* (Aksoy *et al.*, 2013). Specifically, we analyzed this dataset, which included 4584 genotyped individuals (median number of individuals per population: 207 [29–417]), in both BORICE (Koelling *et al.*, 2012) and MLTR (Ritland and Jain, 1981; Ritland, 2002) with default parameters. Because BORICE and MLTR estimates were correlated with each other and BORICE is known to outperform MLTR when maternal genotypes are unavailable (Koelling *et al.*, 2012), we kept only BORICE’s estimates of the family-level outcrossing rate (t) and maternal-line inbreeding coefficients (F) for all subsequent analyses. It is important to note that because removing loci identified as having over 25% null alleles using Micro-Checker (Van Oosterhout *et al.*, 2004) did not significantly impact BORICE’s estimates, we opted for F and t estimates based on all 15 loci (Kuester *et al.*, 2017). We also calculated inbreeding depression (ẟ) using Ritland’s (Ritland, 1990) formula under the assumptions that populations are at inbreeding equilibrium and that the genetic markers used are effectively neutral (Ritland, 1990; Goodwillie *et al.*, 2005).

### Trait correlation

To investigate the degree of correlation between our floral traits and mating system parameters we chose to run separate analyses for our composite floral trait (i.e., ASD) and for our direct floral measurements (TAL, SL, CL, and CW). We made this decision because of the likely cause-effect relationship of ASD with selfing rates (Chang and Rausher, 1998) and the lack of significant correlations between ASD and the other floral traits, albeit the remarkably high correlations among all four direct measurements (Fig. S1). First, we calculated pairwise Pearson’s simple correlation coefficients between ASD and t, F and δ. In addition, we explored the degree of multivariate correlation between our four direct floral measurements (TAL, SL, CL, and CW) and all three mating system parameters (t, F, δ) by running a canonical correlation analysis (CCA) in R3.3.3 (R Core Team, 2017) using package CCA (González and Déjean, 2012). This latter analysis identifies a set of axes that maximize the correlation between two sets of variables (floral and mating system variables in our case) and hence quantifies the extent and significance of their multivariate relationship (Hotelling, 1936). For subsequent analyses we kept the first pair of CCA axes, which as expected shows the strongest correlation.

### Geographic structure

To assess how mating strategies are structured in *I. purpurea* populations over our study area, we investigated how mating system parameters as well as their correlation with floral traits vary based on geographic location. First, we obtained proxies for floral-mating system correlation strength for each population by rerunning the correlation analyses described in the previous section in a leave-one-out manner. That is, we recalculated the Pearson’s and CCA correlation coefficients after removing each population in turn. We then used the difference between the absolute values of the leave-one-out correlation estimate and the all-data correlation estimate as our index of local population correlation strength (λ, hereafter). Positive values of this index (leave-one-out estimate > all-data estimate) indicate a stronger local association than the global association, whereas negative values (leave-one-out estimate < all-data estimate) indicate a weaker local association. Next, we assessed the association of both, individual parameters and λs, with geographic location by running independent multivariate linear regressions for each element or λ against longitude and latitude. In addition, we assessed the degree of global and local spatial autocorrelation on these data by calculating global and local Moran’s I (Moran, 1950; Anselin, 1995) in R3.3.3 (R Core Team, 2017) using packages ape (Paradis, 2011) and spdep (Bivand and Piras, 2015), respectively. For the local analysis, we adjusted p-values for multiple comparisons using a (Benjamini and Hochberg, 1995) false discovery rate method.

### Environmental influence on individual trait variation

To examine the relationship between the 8 selected environmental factors and individual mating system traits, we ran Ordinary Least Squares (OLS) multivariate linear regressions for each mating system trait and performed backward stepwise variable selection using a resampling model calibration strategy with 500 bootstrap replicates. This strategy allows for bias-correction of error estimates based on nonparametric smoothers and hence avoids possible overfitting given our sample size (Harrell, 2015). All these analyses were run on standardized environmental variables using the package rms (Harrell, 2017) in R3.3.3 (R Core Team, 2017). In addition, we estimated the relative contribution of each predictor retained by comparing their associated regression coefficients (i.e., the expected change in the independent variable per unit change of a predictor when all other predictors in the model remain constant). Given our sample size, however, no interactions were included in any model.

Because several OLS assumptions might be violated by our dataset we additionally chose to run homologous model-free regressions using machine-learning tools. Specifically, we opted to run Random Forest (RF) regressions because they deal efficiently with i) the large p-small n problem (large number of predictors relative to observations), ii) non-linear relationships between independent and predictor variables, and iii) predictors multicollinearity (Breiman, 2001; Strobl *et al.*, 2008). RF regressions use an ensemble of multiple-regression trees to fit subsets of the data onto the different predictors by minimizing the sum of square errors and summarize this ensemble of trees by bootstrapped aggregation (a.k.a. bagging; Breiman, 1999). To select the best set of predictors in these regressions, we iteratively fitted the RF algorithm by removing in each iteration the environmental predictor variable with the unscaled smallest variable importance (i.e., backwards variable selection) until the out-of-bag error (i.e., error rate from samples not used in the construction of a given tree) did not decrease any further (Díaz-Uriarte and Alvarez de Andrés, 2006; Strobl *et al.*, 2008). We chose this greedy method of variable selection because an exhaustive search over all possible subsets of predictors is computationally prohibitive. The relative contribution of predictors in the final model was then assessed by the unscaled increase in the mean square error (MSE) of the prediction after removal of each variable (Liaw and Wiener, 2002; Strobl *et al.*, 2008). Finally, the respective functional relationship of each predictor with the independent variable was recovered using partial dependency plots [Citation error], which are graphical visualizations of the marginal effect of a given variable when the other variables are kept constant. All these analyses were run in R3.3.3 (R Core Team, 2017) using package randomForest (Liaw and Wiener, 2002).

### Environmental influence on traits correlation

Finally, we evaluated how the univariate trait correlation was impacted by environmental variation by re-calculating Pearson’s correlation coefficients on subsamples of populations grouped according to their environmental variable values. We followed a similar procedure re-estimating the multivariate trait correlation by re-running the CCA analysis on environmentally grouped sets of populations. Specifically, we independently grouped each environmental variable into quantiles and used these groups to split our set of populations based on similarity on each environmental variable. We then separately calculated the simple and CCA correlation coefficients between floral traits and mating system parameters (as done above) for each subsample. We assessed the significance of the effect of this environmental grouping by comparing the correlation coefficient obtained against similarly obtained coefficients from a set of 100 randomly split datasets that share the same number of observations for each split as the environmentally grouped data.

## RESULTS

### Trait correlation

As previously found (Chang and Rausher, 1998), ASD was significantly correlated with outcrossing rate (t) (Fig. 2). Yet, ASD was not significantly correlated with either inbreeding coefficient of maternal individuals (F) or inbreeding depression (δ) in our dataset. All other floral traits were correlated in a multivariate manner with these three mating system parameters, although the canonical axes themselves were not significant (Table S2). In this latter analysis, corolla (CL) and tallest statement (TAL) lengths, which are themselves significantly correlated with each other (Fig. S1), showed the strongest effect on mating system parameters. With other floral variables held constant, increments in CL were associated primarily with an increase in t and a decrease in δ, whereas increments in TAL had a slightly weaker but opposite effect.

**Figure 2.**
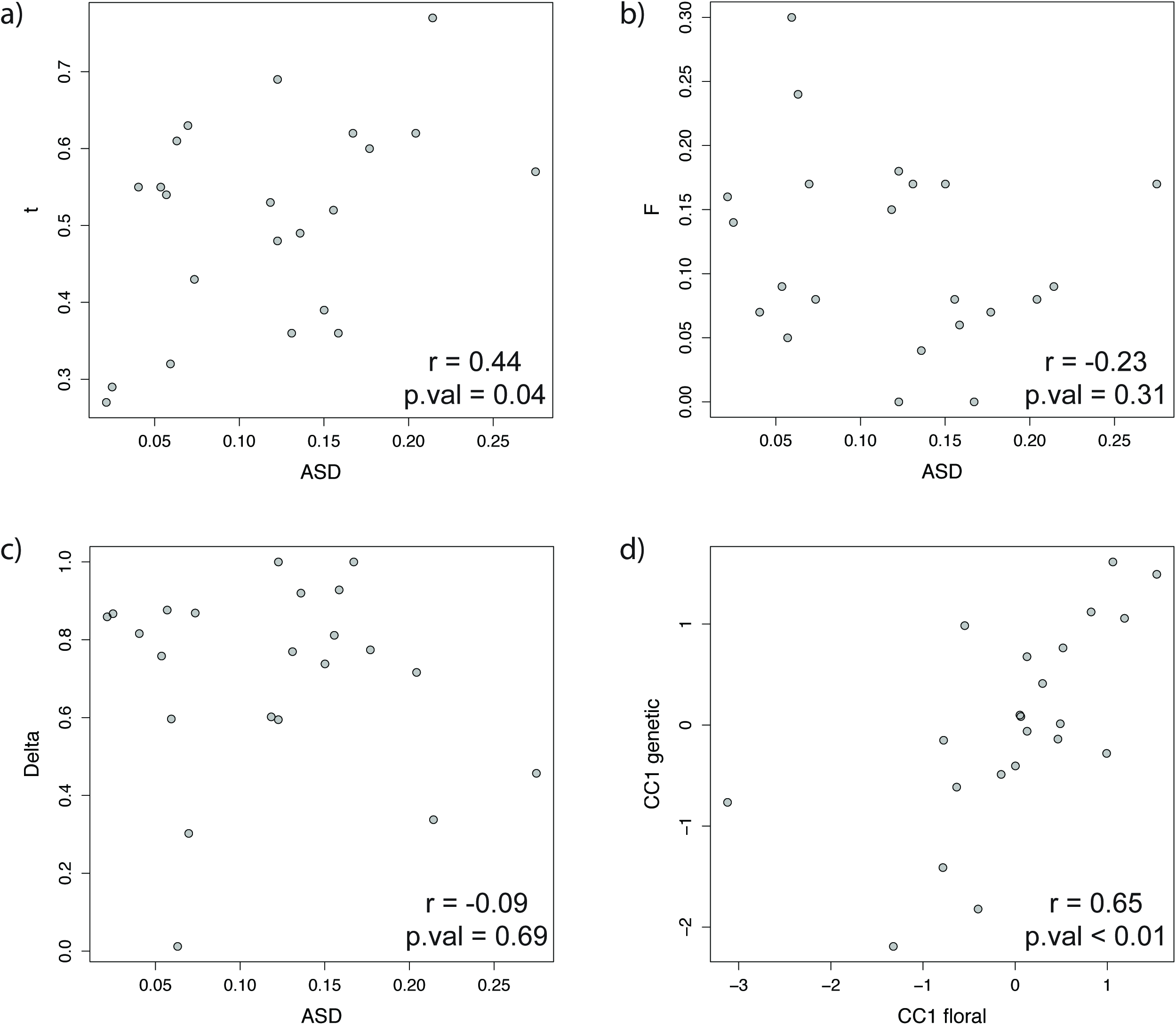
Covariation between floral traits and mating system parameters. a) anther-stigma distance (ASD) and outcrossing rate (t), b) ASD and inbreeding coefficient (F), c) ASD and inbreeding depression (δ), d) first canonical correlation axes of floral and genetic components.

### Geographic structure

We did not uncover significant geographic structure across populations for ASD, for the three examined mating system parameters, or for their correlation with floral traits in *I. purpurea.* No mating system component or floral-mating system correlation index (λ) was significantly associated with either latitude or longitude (Fig. S2), and we did we not find evidence of spatial autocorrelation, as measured by global Moran’s I, in any of our individual variables or λs (Fig. 3). In addition, no population exhibited local spatial autocorrelation in any of our traits (Table 1), meaning there was no evidence for geographically proximate populations showing similar trait values in the study range examined. Further, there was no evidence of local spatial autocorrelation in the correlation indexes (Table 1).

**Figure 3.**
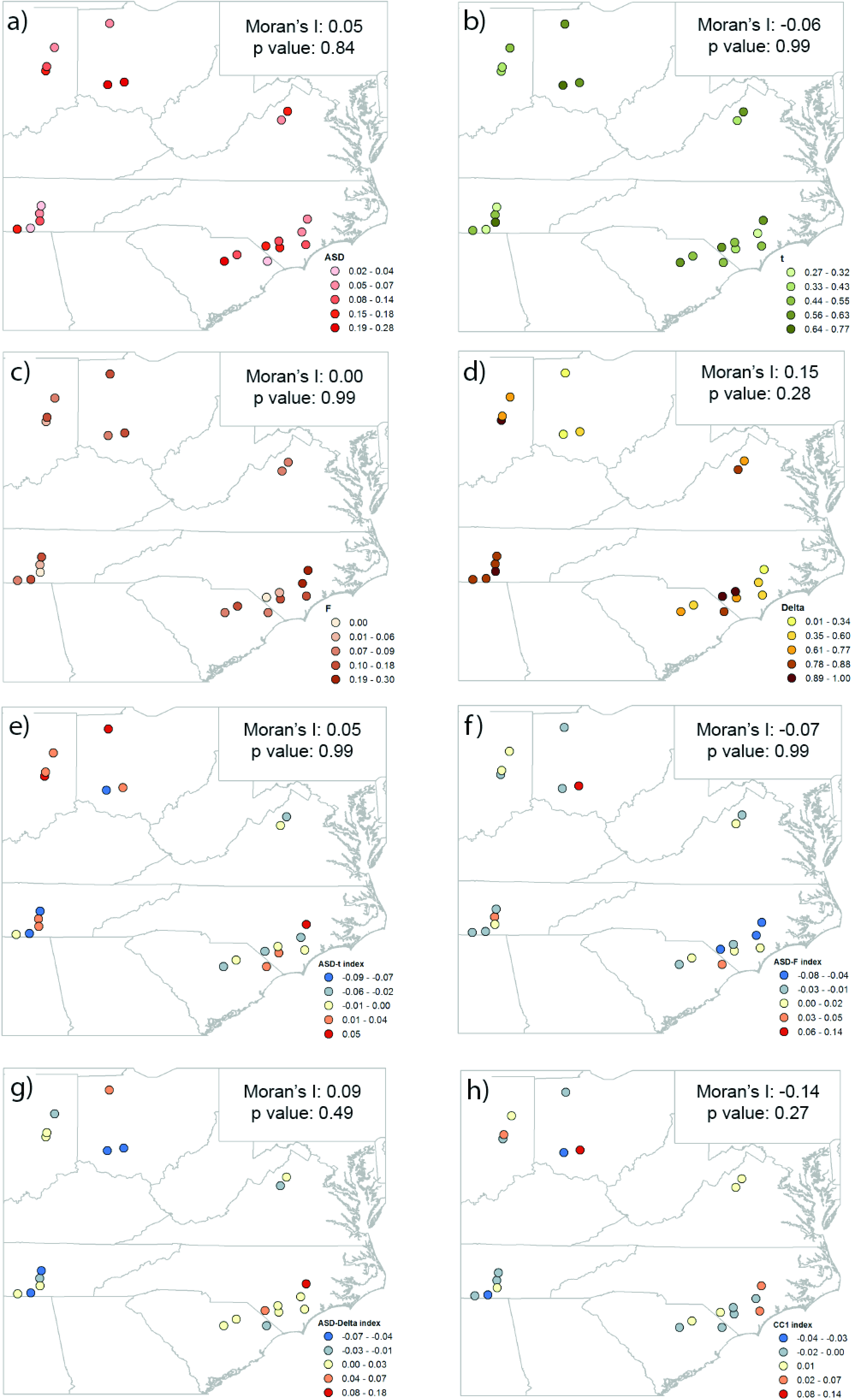
Geographic variability in mating system parameters and their covariation with floral traits. a) anther-stigma distance (ASD), b) outcrossing rate (t), c) inbreeding coefficient (F), d) inbreeding depression (δ), e) ASD-t λ, f) ASD-F λ, g) ASD-δ λ, h) CC1 λ. Global Moran’s I estimates and associated p-values, after correction for multiple testing, are reported.

**Table 1.**
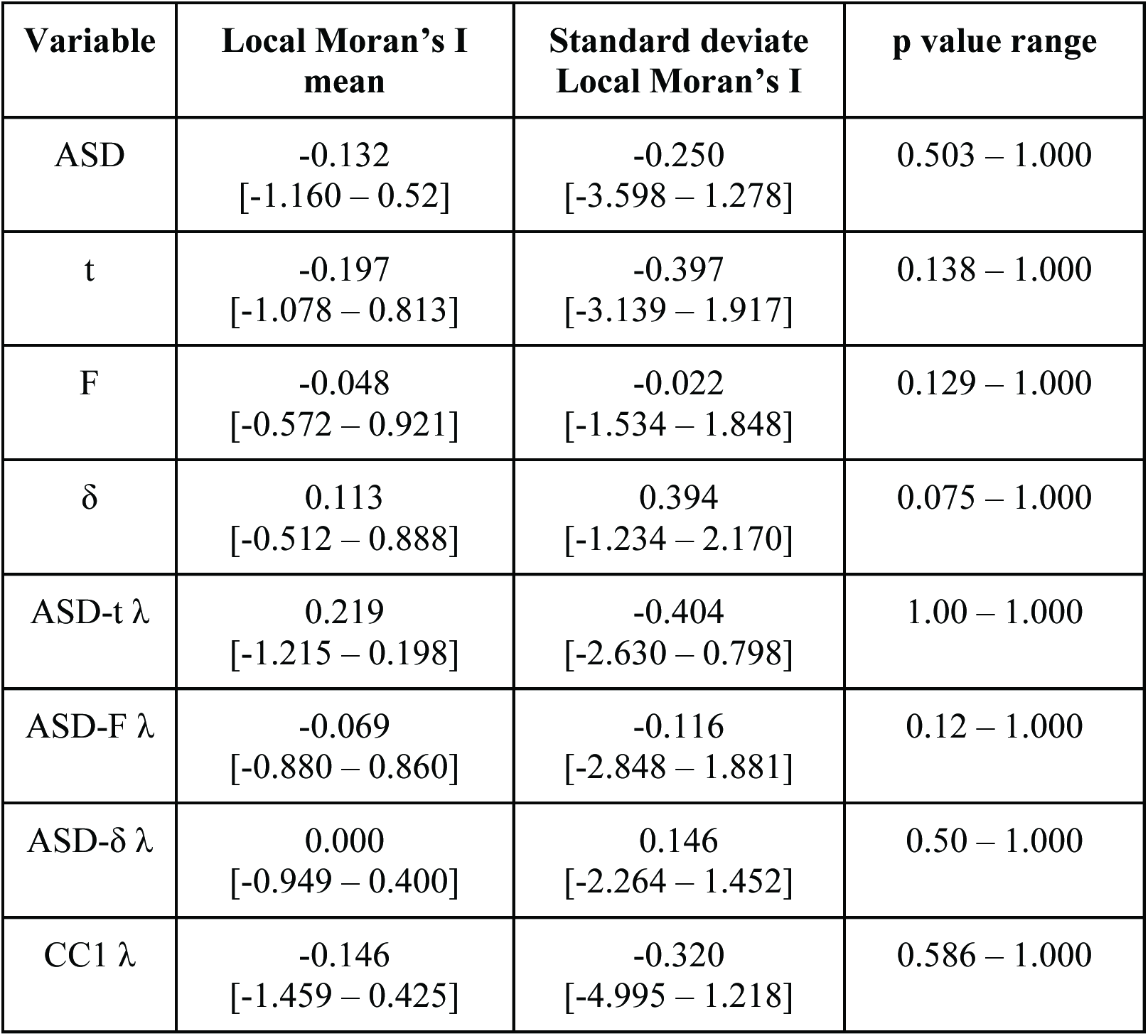
Summary of Local Moran’s I analyses on the extent of local spatial correlation among mating system traits and floral-mating system covariation (λ).

### Environmental influence on individual trait variation

Both OLS and RF regressions identified unique sets of environmental predictors for the different components of *I. purpurea*’s mating system (Tables 2 and 3). Despite the different statistical approaches (Breiman, 2001), resulting in different total number of predictors retained (being in most cases greater in the RF regressions; Tables 2 and 3), both methods robustly identified the same top predictors for all four mating system parameters as well as a relatively congruent relationship between predictors and parameters (Figs. S3 and S4). For example, relative humidity, which was identified as the stronger predictor of ASD under both methods, showed an inversely proportional relationship with ASD in the OLS regression (Fig. S3a) and a thresholded negative association with ASD in the RF regression (Fig. S4a). Across both sets of regressions, we found that herbicide resistance (survival rate after glyphosate application) was the most common predictor (retained in 5 out of 8 regressions), explaining in some cases up to 34% of the variance in mating system traits (Tables 2 and 3). The second most common predictor of mating system traits in our analyses was temperature range (retained in 4 out of 8 regressions), which explained up to 21% of the variance. Nonetheless, the relative importance of these factors—measured as the net effect that change in the predictor causes in the response trait (OLS) or as the increase in the prediction error the predictor removal causes (RF)—vary significantly across mating system traits (Tables 2 and 3). For instance, in both OLS and RF regressions annual temperature range was identified as the most important predictor of inbreeding depression (δ), but only as the third most important predictor of inbreeding coefficient (F). Likewise, the shape of their relationship varies across mating system parameters in an independent manner. For example, whereas the inbreeding coefficient (F) showed a negative relationship with annual temperature range (Figs. S3c and S4c), inbreeding depression (δ) showed a positive association with this environmental factor (Figs. S3d and S4d).

**Table 2.** Summary of Ordinary Least Square linear regressions of mating system parameters on environmental variables. Results obtained after 1000-bootstrapped backward selection are presented. Corresponding simple linear regression results are given in Table S3.

| Dependent | LR<sup>(1)</sup> $\chi^2$ | p value | R<sup>2</sup> | Retained predictors<sup>(2)</sup> | Beta coefficients<sup>(3)</sup> |
| --- | --- | --- | --- | --- | --- |
| ASD | 9.11<br>(3, 18) | 0.028 | 0.34 | Elevation | -0.064<br>(±0.026) |
|  |  |  |  | **Relative humidity** | -0.052<br>(±0.018) |
|  |  |  |  | Mean temperature | -0.049<br>(±0.022) |
| t | 9.05<br>(1, 20) | 0.002 | 0.34 | **Herbicide resistance** | -0.078<br>(±0.024) |
| F | 18.44<br>(4, 17) | 0.001 | 0.57 | **Herbicide resistance** | 0.059<br>(±0.014) |
|  |  |  |  | Annual precipitation | -0.039<br>(±0.018) |
|  |  |  |  | **Temperature range** | -0.038<br>(±0.018) |
|  |  |  |  | Mean Temperature | 0.037<br>(±0.019) |
| Delta | 5.12<br>(1, 20) | 0.024 | 0.21 | **Temperature range** | 0.113<br>(±0.049) |

1. Model likelihood ratio Chi-square statistic. Degrees of freedom are below in parentheses 2. Predictors are ordered by absolute beta coefficient magnitude. Predictors also identified in Random Forest regressions are in bold. 3. Standard errors are in parentheses underneath.

**Table 3.** Summary of Random Forest regressions of mating system parameters on environmental variables. Only results obtained after recursive backward selection are presented.

| **Dependent** | **MSE<sup>(1)</sup>** | **% Variance explained** | **Pseudo-R<sup>2</sup>** | **Retained predictors<sup>(2)</sup>** | **Variable importance<sup>(3)</sup>** |
| --- | --- | --- | --- | --- | --- |
| ASD | 3.67e<sup>-3</sup> | 24.64 | **0.50**<br>(0.02) | **Relative humidity**<br>Annual precipitation<br>Herbicide resistance | 1.18e<sup>-3</sup><br>0.89e<sup>-3</sup><br>0.44e<sup>-3</sup> |
| t | 9.26e<sup>-3</sup> | 46.62 | **0.69**<br>(<0.01) | **Herbicide resistance**<br>Soil coarseness<br>Herbicide rate | 11.77e<sup>-3</sup><br>1.72e<sup>-3</sup><br>0.82e<sup>-3</sup> |
| F | 4.76e<sup>-3</sup> | 11.55 | 0.35<br>(0.11) | **Herbicide resistance**<br>Relative humidity<br>**Temperature range** | 0.88e<sup>-3</sup><br>0.64e<sup>-3</sup><br>0.41e<sup>-3</sup> |
| Delta | 53.96e<sup>-3</sup> | 7.65 | 0.28<br>(0.21) | **Temperature range**<br>Flower size<br>Herbicide use<br>Soil coarseness<br>Mean temperature<br>Elevation<br>Annual precipitation<br>Herbicide resistance | 8.30e<sup>-3</sup><br>5.73e<sup>-3</sup><br>5.48e<sup>-3</sup><br>3.57e<sup>-3</sup><br>3.30e<sup>-3</sup><br>2.34e<sup>-3</sup><br>2.02e<sup>-3</sup><br>1.67e<sup>-3</sup> |

1. Mean of squared errors 2. Predictors are ordered by relative variable importance. Predictors also identified in Ordinary Least Square regressions are in bold. 3. Measured by the decrease in model mean square error after removal of the corresponding variable

In summary, the regressions identified idiosyncratic environmental associations for the four mating system parameters analyzed. ASD decreased as relative humidity increased, elevation and mean temperature increased (OLS only), and that it peaks at intermediate values of annual precipitation and herbicide resistance (RF only) (Figs. S3 and S4). In contrast, as previously observed (Kuester *et al.*, 2017), outcrossing rate is primarily affected by herbicide resistance, being the lowest at higher resistance values (soil coarseness and herbicide use are comparatively minor predictors in RF; Tables 2 and 3). Similarly, the inbreeding coefficient was strongly associated with herbicide resistance and proportionally increased as resistance increased, as previously described (Kuester *et al.*, 2017). The inbreeding coefficient (F) was also positively associated to mean temperature (OLS only) and relative humidity (RF only), and inversely associated to annual precipitation (OLS only) and annual temperature range (OLS and RF) (Tables 2 and 3). On the other hand, inbreeding depression was primarily explained by annual temperature range (Tables 2 and 3) and showed a positive relationship with this environmental factor. It is important to note however, that most other factors analyzed were retained in the RF regression for this latter trait; yet, the overall predictive power of this RF regression was relatively low. Thus, while we found significant associations between components of the mating system and environmental factors, the relationships between these components and environment factors was not congruent.

### Environmental influence on trait correlation

Analyses on how the floral-mating system correlations varied across environmental gradients indicated that no natural environmental factor significantly influenced the univariate correlation between ASD and t, F or δ (results not shown). Nor was there any significant association between natural environmental factors and the multivariate correlation between our floral morphological measurements (TL, SL, CL, and CW) and mating system parameters (results not shown). In contrast, herbicide resistance and herbicide use significantly impacted the correlation between ASD and outcrossing rate and ASD and inbreeding coefficient, respectively. Specifically, the ASD-t correlation was stronger at the greatest herbicide resistance values (Fig. 4a), whereas the ASD-F correlation was weaker at moderately high herbicide use (Fig. 4b). No significant effect of either herbicide use or herbicide resistance on the univariate correlation between ASD and inbreeding coefficient (Fig. 4c) or the multivariate correlation between floral and mating system parameters was recovered (Fig. 4d).

**Figure 4.**
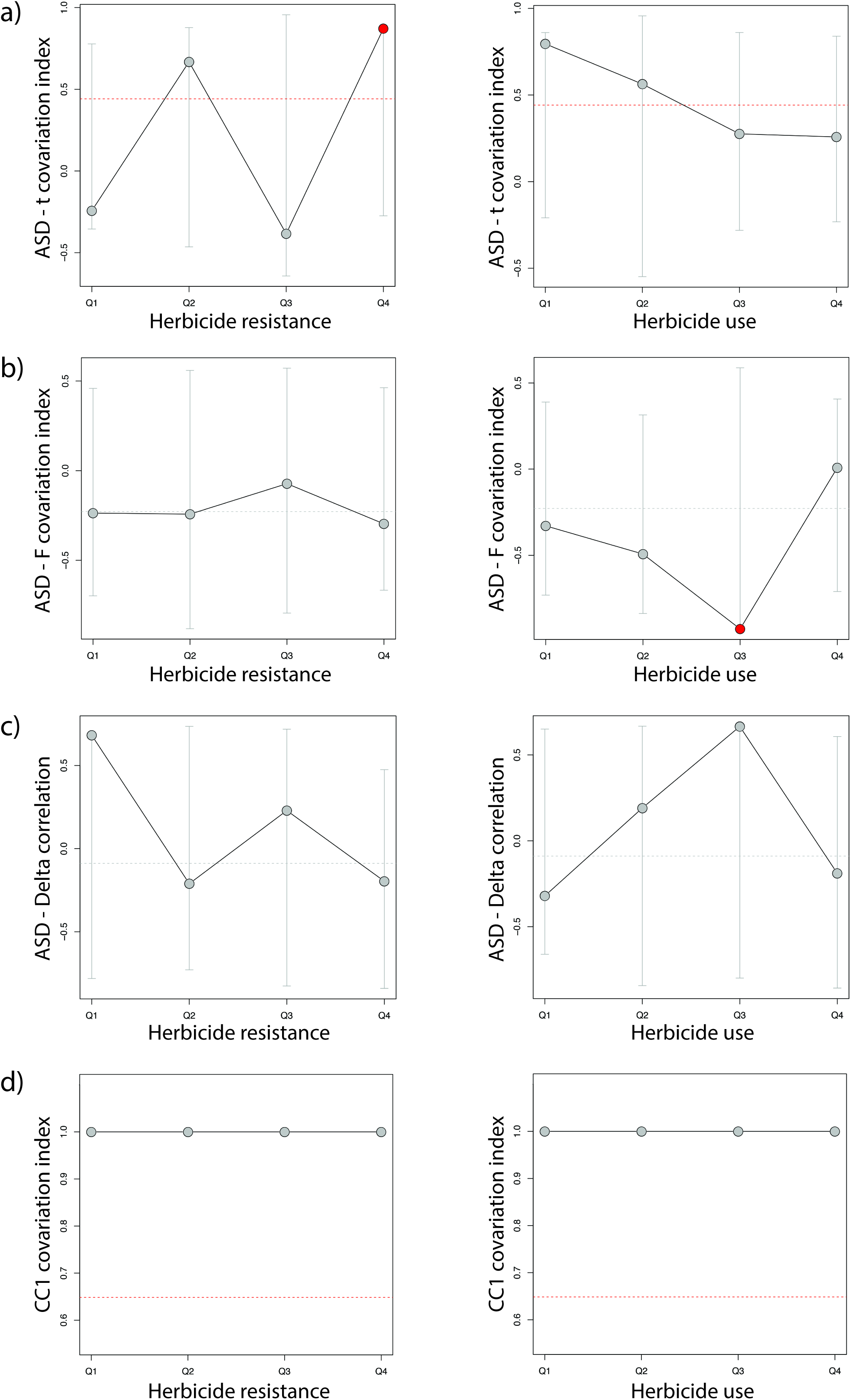
Covariation of floral traits and mating system parameters across anthropogenic selection regimes. Estimates of Pearson’s (a-c) and CCA (d) correlation coefficient across environmentally binned samples are shown by circles, whereas a dashed horizontal line indicates the global estimate. Grey vertical lines indicate 95% confidence estimates based on random resampling. Significant estimates are highlighted by brighter symbols.

## DISCUSSION

Our study identifies disparate environmental factors that influence variation in the mating system of *I. purpurea* across a significant portion of its range. While ASD and the level of inbreeding depression were primarily associated with natural environmental variation, the outcrossing rate and inbreeding coefficient were most strongly associated with the level of herbicide resistance (Kuester *et al.*, 2017). The selection pressure imposed by herbicide use also seems to influence the strength of the association between ASD and selfing and inbreeding. Particularly noteworthy, we did not recover any other environmental influence on the overall weak association between outcrossing rate, inbreeding coefficient, and inbreeding depression with floral traits (TAL, SL, CL, CW, and ASD). Further, we did not find significant geographic structure in any of the mating system parameters explored or their inter-correlation. Taken together, these results suggest that different components of *I. purpurea*’s mating system are presumably under multiple different selective pressures, and that parameters of the mating system and floral traits in this species are not tightly linked. Importantly, these results highlight the complex influence of environmental factors on the mating system of this agricultural weed, and show that human influence is currently a major component of the selfing/outcrossing balance across its populations.

### The complexity of mating systems

Compelling empirical evidence supports an association between individual mating system components and environmental conditions, such as the one found here. For example, outcrossing rate has been found to covary in a variety of plant systems with elevation (Neale and Adams, 1985), humidity (Brown *et al.*, 1978; Shea, 1987), and temperature (Holtsford and Ellstrand, 1992). Similarly, ASD has been found to strongly respond to environmental factors, including humidity (Elle and Hare, 2002; Van Etten and Brunet, 2013), water and nutrient availability (Vallejo-Marín and Barrett, 2009), light regime (Brock and Weinig, 2007), and temperature (Lankinen *et al.*, 2016). Also in agreement with our findings, plenty of studies have identified an association between ASD (herkogamy) and outcrossing rates (Chang and Rausher, 1998; Motten and Stone, 2000; Takebayashi *et al.*, 2006; but see Medrano *et al.*, 2005), and some have identified associations between ASD and inbreeding depression within individual populations (Takebayashi and Delph, 2000; Stone and Motten, 2002; but see Carr *et al.*, 1997). Yet, a relatively small number of studies have simultaneously explored variation patterns of multiple mating system parameters in natural populations across environmental gradients. Among those that have, a variable strength of association is often identified (e.g., Lankinen *et al.*, 2016), which has prompted the hypothesis that variation in the different mating system parameters, such as selfing and inbreeding depression, is more strongly conditioned by other factors (e.g., population size and intraspecific competition for pollinators) than by each other (Johnston and Schoen, 1996; Spigler *et al.*, 2010). Our findings support this hypothesis of relative independence of mating system components (Johnston and Schoen, 1996; Dudley *et al.*, 2007) as well as its expectation of limited cohesive responses among mating system components to environmental variation. The idiosyncratic responses recovered also suggest that mating system determination is very complex and driven by multiple interacting processes. Yet, it remains to be investigated what those interacting processes are and how they are (directly or indirectly) shaped by environmental factors.

Further attesting the complexity of mating system variation is the lack of geographic structure recovered across all mating system parameters. This highlights the importance of fine-tuning mating strategies (through plasticity and/or adaptation) to local environmental conditions. Considering the dramatic evolutionary consequences that reproductive strategies may carry (Kalisz, 1989; Glémin *et al.*, 2006), individuals with reproductive strategies ill-matched to their environmental reality are expected to experience strong detrimental fitness consequences. In line with this expectation, population-level differences in selfing rates are usually associated with habitat quality, as pollen flow is more limited in harsher habitats (Griffin and Willi, 2014; Matos Paggi *et al.*, 2015). More generally, the lack of geographic structure also highlights the importance of local interactions between multiple environmental factors and hence, the complex nature of environments plants have to interact with (Holtsford and Ellstrand, 1992; Sagarin *et al.*, 2006).

At least in *I. purpurea*, there is a combination of multiple different environmental factors acting on different components of its mating system. Inbreeding coefficient, for example, is associated with both climatic variation and herbicide selective pressure (see below). Inbreeding depression is instead mostly explained by annual temperature range, with stronger depression in more temperature-seasonal environments, raising the possibility of additional environmentally driven processes contributing to determine the fitness consequences of inbreeding. For instance, it is possible that the strong association with temperature seasonality, which is a proxy for environmental stability, reflects the more detrimental effect of inbreeding in more stressful environments (Armbruster and Reed, 2005; Cheptou and Donohue, 2011). In contrast, the negative association between ASD and relative humidity might be best explained by selection for reproductive assurance as pollinators’ visitation rates have been found to decrease with relative humidity at least in some plant species (Wang *et al.*, 2009). While these hypotheses definitively require further testing, especially considering the impossibility of assessing direct causality from statistical associations (Mac Nally, 2000), the pattern of differential responses to environmental factors that emerges unequivocally demonstrates the complex integration of mating system strategies and calls for the inclusion of biologically realistic complexity in mixed mating system models.

### The pervasive role of humans

While a significant association between herbicide resistance and selfing rate has been previously uncovered in *I. purpurea* (Kuester et al., 2017), it remained unclear the relative importance of this association in relation to other environmental factors or whether this was a spurious association mediated by other environmental factors (herbicide resistance is itself correlated with precipitation and soil variables; Alvarado-Serrano and Baucom unpublished data). It was also unknown if herbicide resistance similarly impacted other mating system components in this species and hence, what is the overall impact of herbicide usage on its reproductive strategies. Here, by simultaneously exploring the association of mating system components with multiple environmental variables, we provide further support to the hypothesis that anthropogenic-driven selection indeed plays a major role in *I. purpurea*’s mating system dynamics. *Ipomoea purpurea*’s response to the continuous application of glyphosate is by far the strongest predictor of outcrossing and inbreeding rates, and it has a significant impact on the association of these traits and ASD. Specifically, our results suggest that ongoing selection for herbicide resistance in this species may simultaneously favor increased selfing rate and the inbreeding coefficient, promoting a stronger link between them (Fig. 4). This is because the strong selective pressure imposed by herbicide application presumably reduces the number of conspecifics available to mate with (favoring shorter ASD and increased selfing) and also increases the fitness costs of mating with non- or less-resistant plants (favoring inbreeding). Under these circumstances, other (arguably weaker) selective forces normally acting rather independently on different mating system components might be superseded by the remarkably strong selection imposed by herbicide use (Culpepper *et al.*, 2001). In this way, herbicide application is expected to also impact the evolutionary trajectories of weeds by favoring stronger floral integration for increased selfing (Rosas-Guerrero *et al.*, 2011; Fornoni *et al.*, 2016), and by altering the overall genetic constitution of populations (through inbreeding).

Although the impact of human activities on world’s ecosystems is undeniable (Vitousek *et al.*, 1997; Haberl *et al.*, 2007), limited evidence exists of their influence on mixed mating systems in natural populations. The available evidence mostly supports an indirect impact of human activities on mating systems, mediated by habitat modifications (Eckert *et al.*, 2010). Yet, examples of human activities directly conditioning selfing and inbreeding rates and related phenotypic traits remain less common. Our findings not only highlight the extensive cascade of consequences of human activities on species, but also provide further support for humans as direct selective agents of mating strategies (Kuester *et al.*, 2017). While it remains to be seen how prevalent these effects are in less extreme selective regimes, our findings reveal the high evolutionary lability of mating systems and hence its potential sensibility to anthropogenic impacts. Specially considering the relatively short time scale over which it has happened in *I. purpurea* (Duke and Powles, 2008), our results call attention to the need of considering the potential major impact of human-driven selection on such a fundamental life history trait in management and conservation efforts.

### Future directions and conclusions

Although years of research on mixed mating systems have identified a plethora of plausible mechanisms underlying its maintenance (Barrett, 2014), there has been seldom empirical evidence to assess their relative contribution in natural populations. Our unique dataset, which comprises a rare combination of floral measurements and mating system estimates over a significant portion of a species’ range, allows exploring the potential of some of these proposed mechanisms to explain mating strategies. In particular, this unique dataset in combination with a set of robust statistical analyses unequivocally shows that floral traits and mating system estimates do not tightly covary, and seem to respond to different environmental predictors. This finding supports the existence of an intricate network of interactions between mating system components and environmental variation. The uncovering of such complex interactions calls for future studies focused on disentangling the specific nature of the recovered associations to identify direct or indirect causal links between environmental predictors and species’ selfing/outcrossing balance. Yet, even as this work is in progress, the results from the current study offer practical information to forecast plausible consequential responses of species to environmental change, a topic of major importance given the significant evolutionary consequences of variation in mating system patterns.

## ACKNOWLEDGEMENTS

We are grateful to Adam Kuester for collecting the data used in this study, to Megan Van Etten and others in the Baucom lab for helpful discussions and comments on previous drafts. We also thank John Megahan for his illustration of *I. purpurea* flower.

## DATA AVAILABILITY

All original data will be made available in Dryad upon acceptance.

